# Divergent Loop Architecture Shapes Pocket 2 Variation in Shark Legumains

**DOI:** 10.64898/2026.06.04.730232

**Authors:** Micah Eijzenga, Maiya Leibowitz, E. Micheal Henley

## Abstract

Legumain (AEP) is a cysteine protease with a highly conserved catalytic core but variable surface features whose evolutionary and structural diversity remain incompletely understood. To investigate how these features differ across vertebrates, we compared human legumain with six shark orthologs using sequence alignment, AlphaFold based structural modeling, pocket detection, and qualitative docking. All shark sequences retained the canonical His–Cys dyad and β sandwich fold, and structural superposition revealed strong global conservation (RMSD = 0.39 Å). A single surface exposed loop adjacent to a shallow cavity—designated Pocket 2—displayed pronounced sequence divergence. Structural models reproduced this pattern, with Pocket 2 showing the greatest variation in geometry and residue composition across species, including *Callorhinchus milii*, which clustered with elasmobranchs in Pocket 2 features. Although loop conformations were predicted with lower confidence, Pocket 2 was consistently detected in all models and exhibited interspecific differences in volume, shape, and physicochemical environment. Docking of a structurally characterized reference ligand (5KN) into Pocket 2 revealed species specific differences in modeled binding orientations and interaction patterns that were consistent with these geometric variations, though not interpretable as quantitative affinity predictions. Together, these results identify Pocket 2 as a structurally conserved but locally variable region of legumain and highlight it as a candidate site for future experimental and computational investigation. This study is hypothesis generating and provides a framework for examining how localized structural variation may arise within an otherwise conserved protease family.

## Materials/Methods

### Sequence acquisition and curation

Human legumain (AEP) was used as the reference sequence for motif annotation and catalytic domain boundaries (UniProt Q99538). Shark orthologs were identified from publicly available whole genome assemblies at NCBI using the BLAST implementation in Geneious Prime (Kearse et al., 2012). Species were selected based on the availability of high quality genome assemblies and to provide phylogenetic breadth across Chondrichthyes. In particular, *Callorhinchus milii* (Holocephali) was included as an outgroup to the elasmobranch species to broaden evolutionary context. For each species, the top scoring genomic locus was extracted and translated to produce a predicted protein sequence. The nucleotide sequences generated and annotated for this study were deposited under submission SUB16114519 with accession identifiers PZ276389 (*Carcharodon carcharias*), PZ276390 (*Chiloscyllium plagiosum*), PZ276391 (*Hemiscyllium ocellatum*), PZ276392 (*Heptranchias perlo*), PZ276393 (*Heterodontus francisci*), and PZ276394 (*Callorhinchus milii*).

Each predicted protein was aligned to the human reference and manually inspected for the presence of the catalytic His–Cys dyad and the QVVAG motif, which are conserved across vertebrate legumains (Dall & Brandstetter, 2016). Regions corresponding to signal peptides, propeptides, and low confidence terminal extensions were removed so that each curated sequence represented only the catalytic domain. Sequences with frameshifts, premature stop codons, or missing catalytic residues were excluded. The final curated sequences were exported in FASTA format and used for all subsequent analyses.

Multiple sequence alignment was performed with MAFFT using the L INS i strategy (Katoh & Standley, 2013), which is recommended for datasets with conserved cores and variable loop regions. Alignments were inspected manually in Geneious Prime to ensure correct placement of conserved motifs and to avoid misalignment of insertions in surface exposed loops. Residue conservation and per site variability were visualized in Jalview (Waterhouse et al., 2009). This analysis identified a single highly variable region across species corresponding to a surface exposed loop adjacent to a cavity in the human structure. Because this loop displayed both high sequence variability and positional divergence in structural models, we designated the associated cavity as Pocket 2 for focused analysis. The alignment and conservation patterns are presented in Figure 1.

**Figure 1.**
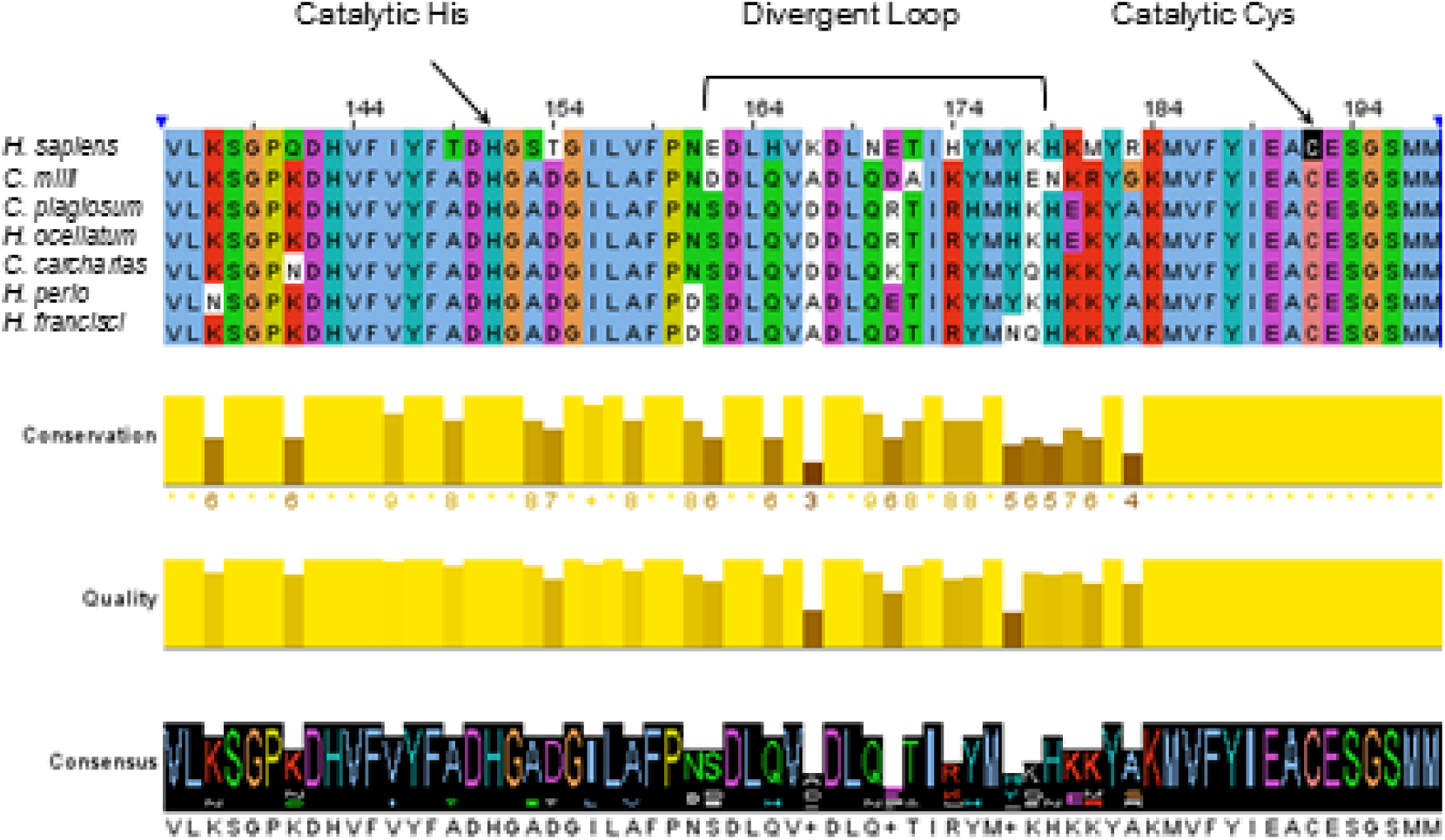
Multiple sequence alignment of human and shark legumain catalytic domains. Alignment of curated catalytic_domain sequences from *H. sapiens* and six shark species. Conserved residues, including the catalytic His and Cys, are highlighted, and a single surface_exposed loop shows pronounced divergence across species. Conservation and alignment_quality scores are shown below the alignment, with a consensus sequence displayed at the bottom.

### Structural modelling and comparative analysis

Structural models for each shark ortholog were generated using ColabFold (Mirdita et al., 2022), which implements the AlphaFold2 architecture (Jumper et al., 2021). For each sequence, five independent models were generated. Representative models were selected based on consistency of loop placement across models and overall structural plausibility. *Carcharodon carcharias* was selected as the representative species for structural comparison with the human reference given the availability of a high-quality whole-genome assembly and its status as a well-characterized elasmobranch. Because the Pocket 2 loop exhibited lower confidence than the catalytic core, pLDDT values for the *C. carcharias* model are provided in Supplementary Figure S1. Because the Pocket 2 loop exhibited lower confidence than the catalytic core, we quantified uncertainty by extracting mean pLDDT values for loop residues and reporting them alongside catalytic core pLDDT values. PAE values for loop to core residue pairs were also summarized to contextualize structural uncertainty.

Representative models were examined in PyMOL (Schrödinger, LLC) to confirm the presence and geometry of the catalytic His–Cys dyad and to verify that the overall β sandwich fold matched the human crystal structure (PDB 4FGU; Dall et al., 2013). Structural comparisons between human and shark models were performed by pairwise superposition of Cα atoms in PyMOL. RMSD and TM scores were calculated for each human–shark pair and are reported in Supplementary Table S1. The RMSD value shown in Figure 2 corresponds specifically to the human–*Carcharodon carcharias* comparison, which exhibited the closest structural similarity among the species examined.

Surface cavities were identified using fpocket (Le Guilloux et al., 2009). All cavities detected in the human structure were mapped onto shark models by structural superposition, allowing identification of equivalent pockets across species. For each pocket, cavity volume, residue composition, and solvent accessibility were recorded. Pocket 2 was consistently detected across all models and incorporated residues from the highly variable loop identified in the sequence alignment. Because the loop was initially highlighted based on sequence variability, we note explicitly that the fpocket screen served as an independent geometric validation of this alignment based hypothesis rather than a blind discovery process.

### Ligand docking and qualitative evaluation

To explore how sequence and structural variation in Pocket 2 might influence ligand accommodation, we extracted the ligand present in the human crystal structure (PDB ligand 5KN) and prepared it for docking using AutoDockTools (Morris et al., 2009). Both ligand and receptor structures were prepared by adding polar hydrogens and assigning Gasteiger charges, and the ligand geometry was energy minimized prior to docking. Docking was performed with AutoDock Vina (Trott & Olson, 2010) using an exhaustiveness of 32, twenty output poses, an energy range of 4 kcal/mol, and a fixed random seed of 2025. Ten independent docking replicates were performed for each species. A cubic grid box was centered on Pocket 2 and sized to encompass the full cavity and adjacent loop region, and the same grid parameters were applied to all species to ensure comparability.

Docked poses were visualized in PyMOL and interaction diagrams were generated with LigPlot+ (Laskowski & Swindells, 2011). Because Vina scores are not directly comparable across structurally distinct binding sites, docking results were interpreted qualitatively, with emphasis on pose stability, interaction patterns, and geometric complementarity rather than quantitative affinity predictions. Statistical comparisons of score distributions were used only to summarize replicate variability and are reported with effect sizes and Dunn’s post hoc tests where appropriate.

### Quality control and uncertainty assessment

Quality control checks were applied throughout the workflow to ensure that conclusions were not driven by single anomalous models or alignments.

These included inspection of alignment columns corresponding to pocket residues, comparison of Pocket 2 geometry across all five ColabFold models per species, extraction of pLDDT and PAE values for loop regions contributing to Pocket 2, and verification that the catalytic dyad geometry was preserved in all representative models. Docking replicates were examined for pose consistency and for whether top poses contacted conserved catalytic residues or variable loop residues. Interpretation of structural differences and docking results was guided by model confidence metrics, and observations dependent primarily on low confidence loop positions were treated as hypothesis generating rather than definitive. All analyses were performed with software versions current as of March 2025.

## Results

### Sequence conservation and divergence across shark legumains

Alignment of the curated catalytic domains showed that all shark orthologs retained the canonical His–Cys catalytic dyad and the QVVAG motif characteristic of vertebrate legumains (Dall & Brandstetter, 2016). Despite this conserved catalytic core, substantial sequence variation was concentrated in a single surface exposed loop (Figure 1). This region displayed elevated amino acid variability and several lineage specific insertions and deletions, whereas residues forming the catalytic cleft and the β sandwich scaffold were strongly conserved across species.

Per site conservation scores calculated in Jalview highlighted a sharp boundary between the conserved catalytic core and the variable loop. In the human structure, this loop borders a shallow surface cavity. Because this region showed the strongest divergence at the sequence level, we considered it a candidate for structural and functional variation among shark legumains, motivating the comparative structural analyses described below.

### Structural conservation of the catalytic fold and divergence in the Pocket 2 loop

Structural models generated with ColabFold (Mirdita et al., 2022; Jumper et al., 2021) reproduced the expected β sandwich architecture of legumain across all shark species. The catalytic His–Cys dyad was positioned consistently in all models, and the overall fold aligned closely with the human crystal structure (PDB 4FGU; Dall et al., 2013). Structural comparison using *C. carcharias* as the representative shark ortholog yielded a global Cα RMSD of 0.39 Å, and an RMSD of 0.56 Å when restricted to residues forming the catalytic site, indicating strong conservation of both the global fold and the active site geometry (Figure 2).

**Figure 2.**
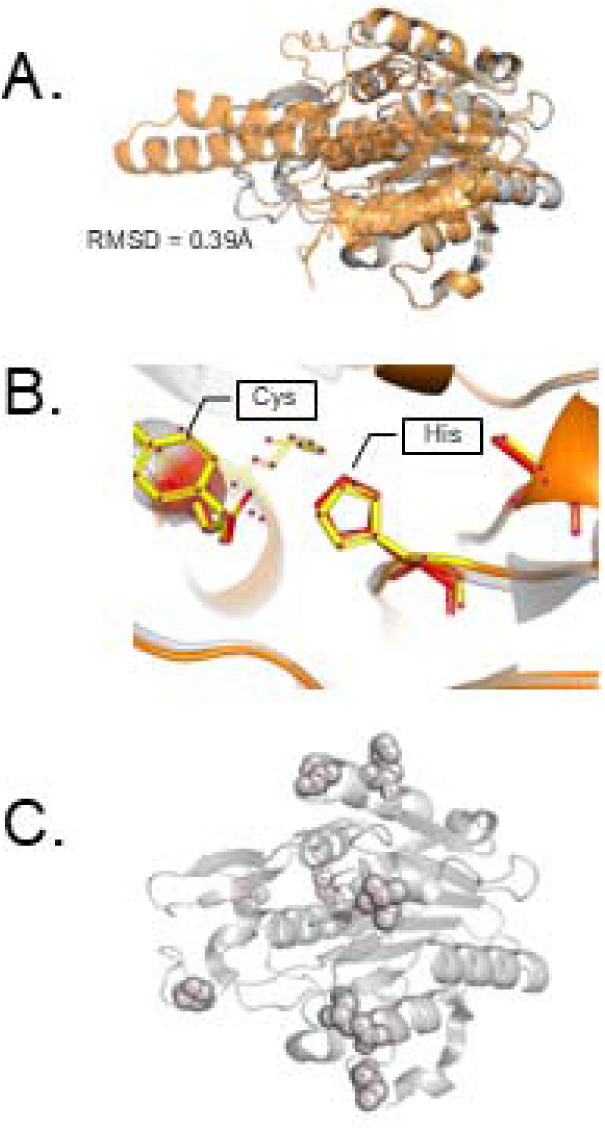
Structural comparison of human and shark legumain models. (A) Superposition of representative human and shark legumain structures showing overall fold conservation (RMSD = 0.39 Å). (B) Close_up view of the catalytic His–Cys dyad, which is preserved across species. (C) Distribution of predicted ligand_accessible regions mapped onto the structural model.

Despite this overall similarity, the loop identified as highly variable in the sequence alignment showed the greatest positional divergence in the structural models. In some species, the loop extended outward relative to the human structure, whereas in others it folded inward toward the cavity. These differences were accompanied by elevated PAE values in the same region, consistent with increased conformational flexibility. Although the precise loop conformations should be interpreted cautiously due to lower pLDDT values in this region (Supplementary Figure S1), the consistent interspecific differences across independent models suggest that the underlying sequence divergence contributes to variation in the geometry of the adjacent cavity, Pocket 2.

### Pocket 2 is conserved as a cavity but varies substantially in geometry and residue composition

Cavity detection using fpocket (Le Guilloux et al., 2009) identified Pocket 2 in all shark models as well as in the human structure. Although the pocket was conserved in position, it exhibited notable interspecific variation in volume, shape, and residue composition (Figure 3). Pocket volumes ranged from compact and shallow in *Heterodontus francisci* to substantially expanded in *Carcharodon carcharias* and *Heptranchias perlo*. These differences were driven primarily by substitutions within the variable loop, including changes in side chain size, polarity, and charge.

**Figure 3.**
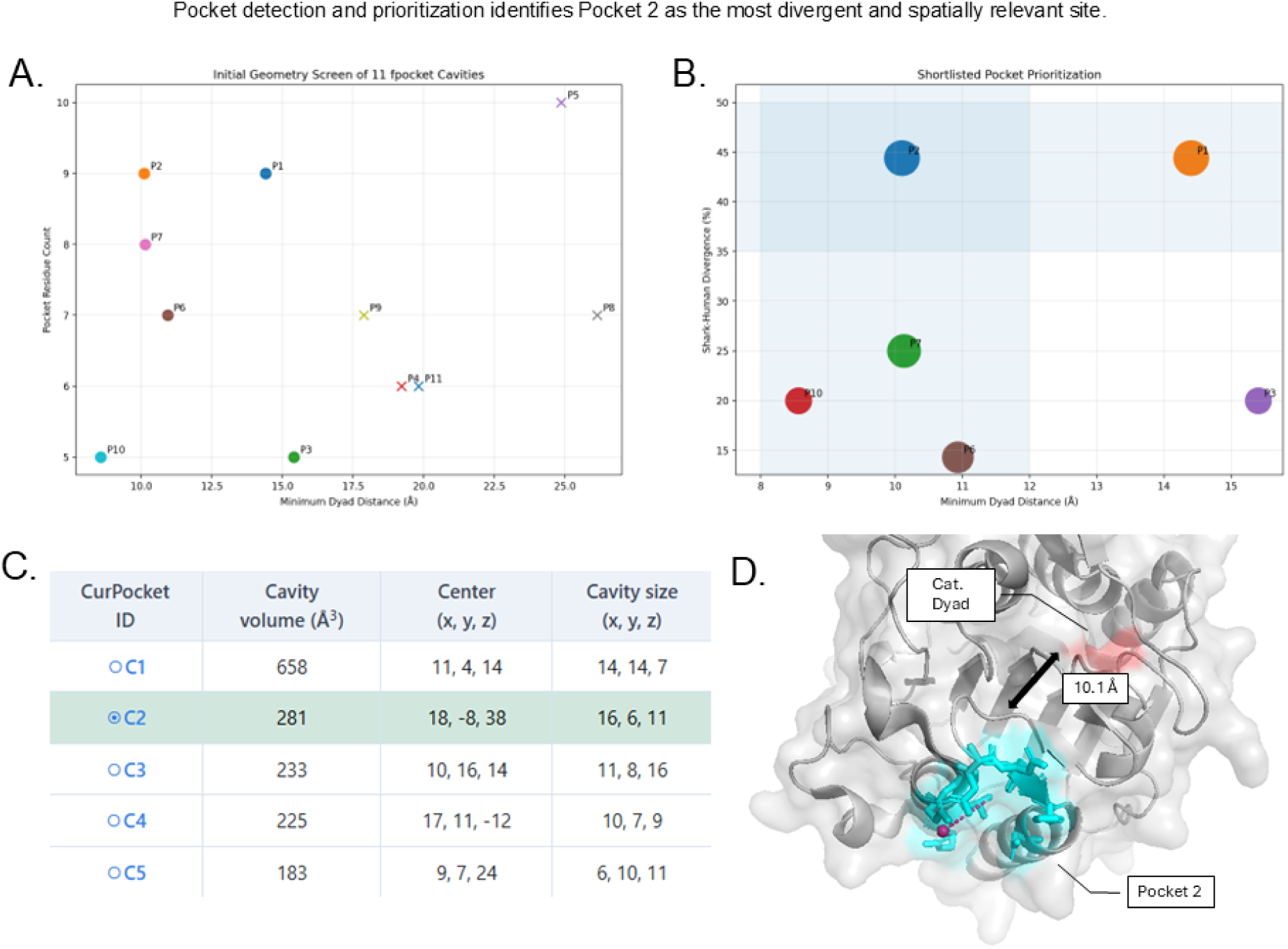
Identification and prioritization of surface cavities, highlighting Pocket 2. (A) Initial fpocket screen of 11 cavities plotted by minimum distance to the catalytic dyad and residue count. (B) Shortlisted pockets ranked by dyad distance and substructure divergence. (C) Cavity properties for the top candidates, with Pocket_2 highlighted. (D) Structural rendering showing the spatial relationship between Pocket_2 and the catalytic dyad (10.1 Å separation).

Residue composition analysis showed that species with larger Pocket 2 volumes tended to possess more polar or charged residues lining the cavity, whereas species with smaller pockets retained predominantly hydrophobic residues. These compositional differences altered the predicted physicochemical environment of the pocket. Although the spatial relationship between Pocket 2 and the catalytic dyad remained conserved, the local variation in geometry and residue identity suggests that Pocket 2 may represent a structurally flexible region whose features differ across species. *Callorhinchus milii*, despite its position as a holocephalan outgroup, clustered with the elasmobranch species in Pocket 2 geometry and residue composition, indicating that the divergence from the human Pocket 2 environment likely predates the holocephalan–elasmobranch split.

### Ligand docking reveals species specific differences in binding orientation and interaction patterns

Docking of the human crystallographic ligand 5KN into Pocket 2 produced distinct binding modes across species (Figure 4). In the human structure, the ligand adopted a pose consistent with its crystallographic orientation, forming hydrogen bonds with residues adjacent to the catalytic dyad. Similar poses were observed in several shark models whose Pocket 2 geometries most closely resembled the human cavity.

**Figure 4.**
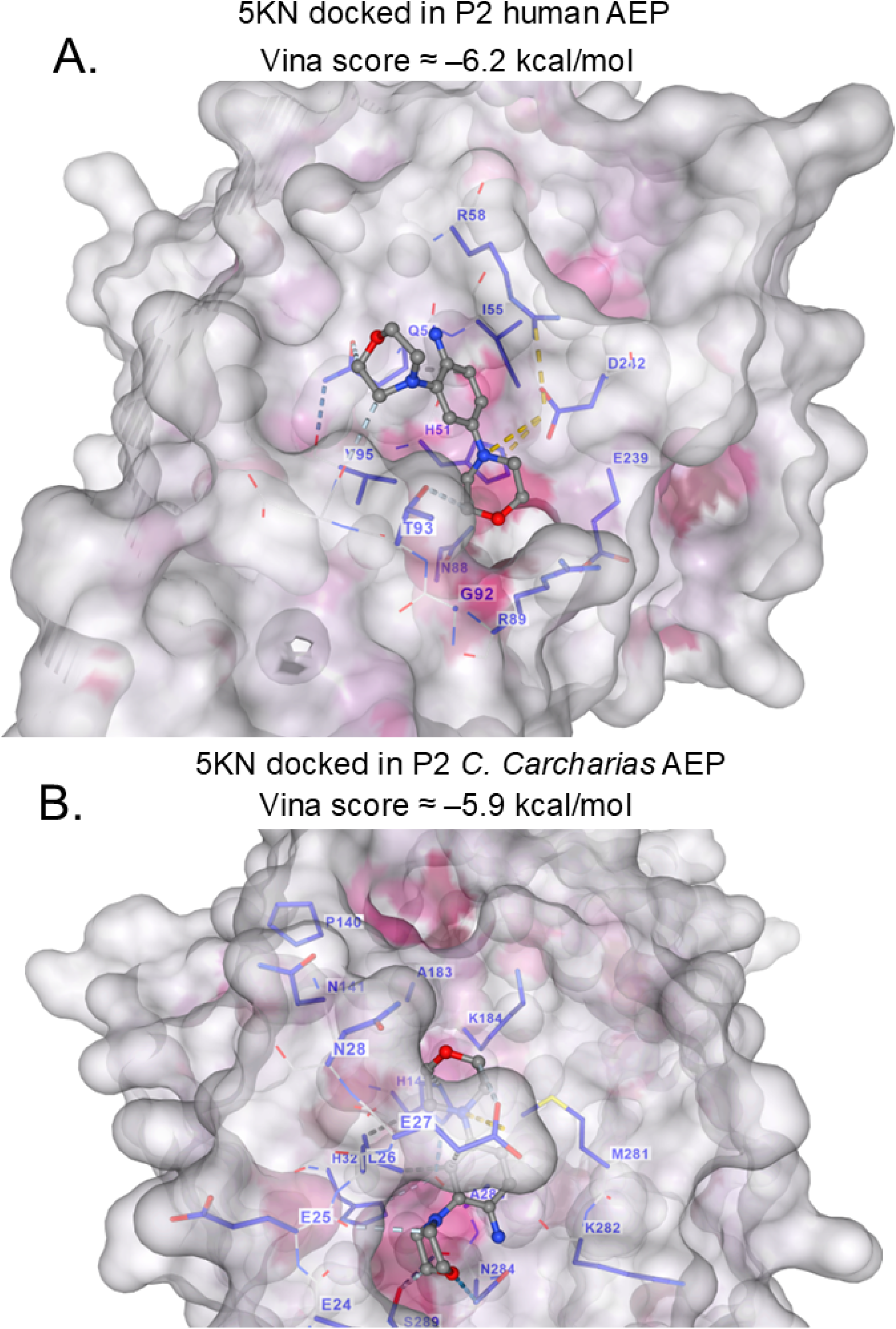
Docking of ligand 5KN into Pocket 2 of human and *C. carcharias* legumain. (A) Docked pose of 5KN in human legumain Pocket 2 (Vina score ≈ –6.2 kcal/mol), with interacting residues labeled. (B) Docked pose of 5KN in *C. carcharias* Pocket 2 (Vina score ≈ –5.9 kcal/mol), showing altered pocket architecture and interaction patterns relative to the human structure.

In species with expanded or reshaped pockets, the ligand adopted alternative orientations. For example, in *Carcharodon carcharias*, the ligand rotated deeper into the cavity and formed additional hydrophobic contacts with loop residues not present in the human structure. In *Heterodontus francisci*, the smaller pocket restricted ligand entry, resulting in a shallower pose with fewer stabilizing interactions. LigPlot+ diagrams highlighted these differences, showing species specific patterns of hydrogen bonding and hydrophobic contacts.

Predicted Vina scores varied across species; however, because Vina scores are not directly comparable across structurally distinct binding sites, these values were used only to confirm that replicate poses were internally consistent within each species rather than to support quantitative cross species comparisons. We therefore interpret the docking results qualitatively, focusing on pose stability, recurrent interaction patterns, and the extent to which modeled ligand orientations were compatible with the observed Pocket 2 geometries. Contact frequency analysis across docking replicates further illustrated that residues within the variable loop contributed disproportionately to ligand interactions in species with expanded pockets, whereas interactions in species with smaller pockets were dominated by conserved residues near the catalytic dyad.

### Structural divergence in Pocket 2 corresponds to differences in modeled ligand accommodation

Comparisons of structural metrics and docking results revealed a consistent relationship between Pocket 2 geometry and the qualitative features of ligand accommodation. Species with the greatest loop divergence also showed the largest deviations in pocket volume and shape, which corresponded to differences in ligand orientation and interaction patterns. These observations suggest that variation in the Pocket 2 loop may influence how small molecules are positioned within the cavity, although experimental validation would be required to determine whether these differences have functional consequences.

Importantly, these conclusions were evaluated in the context of model confidence. Although the Pocket 2 loop exhibited lower pLDDT scores than the catalytic core, the overall pattern of interspecific differences was consistent across independent ColabFold models. Docking results were also stable across replicates, indicating that the observed trends reflect reproducible geometric features of the models rather than stochastic variation in ligand placement. Because loop conformations are predicted with lower confidence, these findings should be considered hypothesis generating and serve as a basis for future experimental or computational validation.

## Discussion

The comparative analyses presented here reveal a consistent pattern in which the catalytic core of legumain is strongly conserved across sharks, while a single surface exposed loop and its associated cavity, Pocket 2, exhibit substantial interspecific variation. The retention of the His–Cys dyad, the QVVAG motif, and the β sandwich scaffold across all species underscores the deep evolutionary conservation of the catalytic machinery. In contrast, the variable loop adjacent to Pocket 2 shows elevated sequence divergence, multiple insertions and deletions, and the greatest positional variability in structural models. These observations suggest that this region represents a flexible and evolutionarily dynamic feature of the legumain fold.

Although the global architecture of shark legumains closely resembles the human structure, the Pocket 2 loop displayed the highest predicted structural uncertainty, as reflected in lower pLDDT values and elevated PAE scores.

The pLDDT visualization for the *C. carcharias* model (Supplementary Figure S1) illustrates this confidence gradient, and loop conformations should therefore be interpreted cautiously. However, the overall pattern of interspecific differences was consistent across independent ColabFold models, suggesting that the underlying sequence divergence contributes to genuine variation in the local structural environment. The persistence of Pocket 2 across species, despite differences in loop orientation and residue composition, further supports the idea that this cavity is a conserved structural element with variable local features. Notably, Callorhinchus milii clustered with the elasmobranch species in Pocket 2 geometry, indicating that the divergence from the human Pocket 2 environment likely predates the holocephalan–elasmobranch split.

The fpocket analysis provided an independent geometric perspective on this variation. Pocket 2 was detected in all species, but its volume, shape, and residue identity differed substantially. These differences were driven primarily by substitutions within the variable loop, which altered the physicochemical properties of the cavity. Species with larger pockets tended to possess more polar or charged residues lining the cavity, whereas species with smaller pockets retained predominantly hydrophobic residues. Although these observations do not establish functional consequences, they highlight Pocket 2 as a region where structural divergence is concentrated.

Docking of the human crystallographic ligand 5KN into Pocket 2 further illustrated how these structural differences may influence ligand accommodation. Because Vina scores are not directly comparable across structurally distinct binding sites, the docking results were interpreted qualitatively. Differences in ligand orientation, depth of insertion, and interaction patterns across species were consistent with the observed variation in pocket geometry. In species with expanded cavities, the ligand adopted deeper or rotated poses that engaged variable loop residues, whereas in species with smaller pockets, ligand placement was more restricted and relied primarily on conserved residues near the catalytic core. In this context, 5KN serves as a structurally characterized reference ligand rather than a biologically validated shark substrate, providing a consistent scaffold for comparing geometric compatibility across species. These modeled interactions should therefore be viewed as indicators of structural accommodation rather than predictions of binding affinity or biological relevance.

The focus on Pocket 2 emerged from an alignment guided hypothesis: the loop adjacent to this cavity was the most variable region across species, and we therefore examined whether the associated cavity also varied. The fpocket screen of all cavities served as an independent validation of this hypothesis rather than a blind discovery process. This approach is common in comparative structural analyses but warrants explicit acknowledgment to avoid the appearance of circular reasoning. The consistent detection of Pocket 2 across species, combined with its pronounced geometric divergence, supports the idea that this region represents a structurally flexible site within an otherwise conserved fold.

Several limitations should be considered when interpreting these findings. First, loop regions are predicted with lower confidence than the catalytic core, and the precise conformations of flexible loops cannot be inferred with high certainty from AlphaFold based models alone. Second, docking results depend on receptor conformation and scoring function assumptions, and therefore provide only qualitative insight into potential ligand accommodation. Third, the biological relevance of the 5KN ligand to shark legumain is unknown; in this study, it serves as a structural probe rather than a functional substrate. Finally, distinguishing adaptive divergence from neutral variation would require explicit evolutionary analyses, which are beyond the scope of this work.

Despite these limitations, the convergence of sequence, structural, and qualitative docking observations highlights Pocket 2 as a region of notable interspecific variation. The consistent divergence of this cavity across sharks suggests that it may represent a lineage variable feature of the legumain fold. These results are best interpreted as hypothesis generating and provide a foundation for future experimental or computational studies aimed at determining whether variation in Pocket 2 influences substrate recognition, regulatory interactions, or other aspects of legumain function.

## Conclusion

This comparative analysis of human and shark legumains identifies Pocket 2 as a structurally conserved but locally variable cavity positioned adjacent to the most divergent loop in the catalytic domain. Although the catalytic machinery and overall β sandwich fold are strongly conserved across species, the Pocket 2 loop exhibits pronounced sequence variation, lower predicted structural confidence, and corresponding differences in modeled cavity geometry. Qualitative docking analyses further suggest that these structural differences may influence how small molecules are accommodated within the pocket, although the biological relevance of these modeled interactions remains to be determined.

Because loop regions are predicted with lower confidence and docking results depend on receptor conformation, the observations presented here should be interpreted as hypothesis generating rather than definitive. Nonetheless, the convergence of sequence divergence, geometric variation, and modeled interaction differences highlights Pocket 2 as a candidate site for future experimental and computational investigation. These findings provide a framework for exploring how localized structural variability may contribute to functional diversity within an otherwise highly conserved protease family.

